# Comparing DNA Extraction and 16s Amplification Methods for Plant-Associated Bacterial Communities

**DOI:** 10.1101/2020.07.23.217901

**Authors:** Cecelia Giangacomo, Mohsen Mohseni, Lynsey Kovar, Jason G. Wallace

**Affiliations:** Department of Biology, University of Georgia, Athens, Georgia; Center for Applied Genetic Technologies, University of Georgia, Athens, Georgia; Institute of Bioinformatics, University of Georgia, Athens, Georgia; Department of Crop & Soil Sciences, University of Georgia, Athens, Georgia

**Keywords:** Amplicon sequencing, Plant microbiome, 16s rRNA, peptide nucleic acid (PNA), blocking oligos, chloroplast, mitochondria

## Abstract

Plant-associated microbes play important roles in global ecology and agriculture. The most common method to profile these microbial communities is amplicon sequencing of the bacterial 16s rRNA gene. Both the DNA extraction and PCR amplification steps of this process are subject to bias, especially since the latter requires some way to exclude DNA from plant organelles, which would otherwise dominate the sample. We compared several common DNA extraction kits and 16s rRNA amplification protocols to determine the relative biases of each and to make recommendations for plant microbial researchers. For DNA extraction, we found that, as expected, kits optimized for soil were the best for soil, though each still included a distinct “fingerprint” of its own biases. Plant samples were less clear, with different species having different “best” options. For 16s amplification, we find that using peptide nucleic acid (PNA) clamps provides the least taxonomic distortion, while chloroplast-discriminating primers are easy and inexpensive but present significant bias in the results. We do not recommend blocking oligos, as they involved a more complex protocol and showed significant taxonomic bias in the results. Further methods development will hopefully result in protocols that are even more reliable and less biased.

## Introduction

Plant-associated microbial communities (the plant “microbiome”) play a significant role in global ecology and agriculture. These microbes affect how plants grow, acquire nutrients, defend against pests and disease, and otherwise carry out many essential activities (reviewed in (Müller et al. 2016; Compant et al. 2019)). There is currently great interest in understanding how plants and microbes interact and how these interactions could be harnessed to improve human activities (Bell et al. 2019; Reid and Greene 2013).

Although the plant microbiome consists of bacteria, archaea, fungi, oomycetes, protists, and other microscopic organisms, most current research focuses on bacteria, followed by fungi. The most common method to profile the bacteria in plant-associated samples is targeted sequencing of the 16s ribosomal RNA gene. This process consists of extracting DNA from plant-associated communities, using PCR to amplify a specific gene sequence (in this case, 16s rRNA) from the bacterial community, sequencing the resulting amplicon, and performing bioinformatic analyses to identify patterns of community composition. The two major processing steps in this method--DNA extraction and PCR amplification--are known to be subject to bias based on the specific methods and conditions used (Pollock et al. 2018). Although no method is completely free of bias, researchers understandably want to use the methods with as little bias as possible.

Bias during DNA extraction comes from incomplete recovery of DNA from the sample. This can be due to microbial cells that are not lysed at equal efficiency or due to chemical inhibitors present in the sample (Pollock et al. 2018). Although phase-separation methods (like CTAB extraction) can be used to recover plant-associated microbial DNA, many researchers choose to use commercial column-or bead-based purification kits due to their greater convenience and consistency. However, these kits have generally been optimized either for the plant itself or for microbes in other environments (soil, feces, etc.). Extracting plant-associated microbes includes the combined challenges of lysing a diverse array of microbes while also dealing with potential chemical inhibitors from the plant itself, and it is unknown how well different kits perform under these circumstances.

Bias during PCR amplification comes primarily from PCR primers and bias from in the DNA polymerase itself (meaning, the enzyme does not amplify all targets equally well). Plant-associated microbiome samples pose an additional challenge in that the most common primers to amplify microbial DNA also amplify plant organelles (chloroplasts and mitochondria). Since organelles are generally much more abundant than microbes in plant tissue, most sequencing reads come from host DNA unless something is done to specifically exclude them. Several methods have been used to selectively discriminate against host DNA in plant-associated samples:

<u>Discriminating primers</u> are designed to selectively bind to bacterial DNA but not plant organellar DNA. Such primers are extremely simple and have been successfully used in Arabidopsis (Bodenhausen, Horton, and Bergelson 2013), poplar (Beckers et al. 2016), maize (Chelius and Triplett 2001; Wallace et al. 2018; Wasimuddin et al. 2019), apple (Shade, McManus, and Handelsman 2013), algae (Thomas et al. 2020), various non-model plants (Vannier et al. 2018; Massoni et al. 2020), and even for the gut microbes of plant-eating insects (Hanshew et al. 2013). The major drawback of discriminating primers is that they cause bias by also discriminating against legitimate bacterial targets (Thomas et al. 2020). Beckers et al. (2016) compared the efficiency of different discriminating primer pairs in poplar and found that the 799F-1391R pair was most suitable for broad taxonomic coverage, followed by 799F-1193R. (The pair used in the current study--799F-1115R--was not tested but is likely similar to these two since the chloroplast discrimination comes from the 799F primer.)

<u>PCR clamps</u> consist of modified oligonucleotides that bind to template DNA during PCR reaction and physically prevent amplification of some products (in this case, organellar DNA). This inhibition usually works by blocking primer sites or preventing the DNA polymerase from physically moving along the template. The most common PCR clamps are peptide nucleic acids (PNAs) (Lundberg et al. 2013; Sakai and Ikenaga 2013), which have been used in a wide variety of plant species (Blaustein et al. 2017; Tkacz et al. 2020; Fitzpatrick et al. 2018). The efficiency of PNA clamps can vary widely across plant species even within similar clades (Fitzpatrick et al. 2018). Researchers have also used locked nucleic acids (LNAs) to similar effect (Ikenaga et al. 2018; Ikenaga and Sakai 2014). LNAs have been used in wheat, soybean, and potato (Ikenaga et al. 2016), but otherwise seem to have limited use in the plant microbiome field.

<u>Blocking oligos</u> are a recently proposed method that uses a nested PCR reaction to “poison” organellar amplicons so that they cannot be used for library preparation (Agler et al. 2016). Because the nested PCR reaction targets DNA specific to the plant of interest, blocking oligos must be custom designed for each target species. In theory they should be compatible among closely related species, but to our knowledge this has not been tested.

To provide guidelines for choosing among these various options, we compared several common methods for DNA extraction and PCR amplification of plant-associated bacteria to identify the potential biases present in them. Although these methods have been used for several years, there has not yet been a head-to-head comparison among them to help researchers make informed choices as to which to use for plants. (Similar comparisons have been done for other species, especially humans, e.g., (Teng et al. 2018; Fiedorová et al. 2019)). Although we did not include analysis of fungal (and other) communities in this study, we acknowledge that they are also important to the plant microbiome and encourage similar studies to be done for them.

## Materials & Methods

### Sample collection

Samples for extraction methods (Experiment I) consisted of leaf tissue from *Arabidopsis thaliana*, corn (*Zea mays*), and soybean (*Glycine max*) growing at the Center for Applied Genetic Technology (CAGT) at the University of Georgia. All plants were grown under standard conditions (growth chamber or greenhouse) with no special treatments or sterilization. *Arabidopsis* samples consisted of 1 mature leaf. For maize and soybean, either 1 or 10 leaf discs were collected using a standard hole punch (~ 8mm diameter); the punch was wiped clean with alcohol between collections to remove any carryover. Soil samples were collected from areas with low (Soil 1) or high (Soil 2) organic matter.

Samples for PCR primer tests (Experiment II) were handled similarly, save that soybean and corn samples came from the Iron Horse Plant Sciences Farm in Watkinsville, Georgia. Two different soil samples (a high-organic landscaped soil and a low-organic native clay soil) were collected around the Center for Applied Genetic Technologies. Defined community DNA (ATCC 20-Strain Even Mix Genomic Material; ATCC MSA-1002) was also tested as a positive control to check for community distortion.

All samples were frozen at −80 C until extraction. Plant tissue was frozen in tubes pre-loaded with two 4.5 mm steel beads for later grinding. Soil was weighed into 200-250 mg aliquots before DNA extraction.

### DNA Extraction

DNA extraction protocols followed the manufacturer's guidelines with minor modifications (e.g., adding small amounts of lysis buffer before grinding because it improved grinding efficiency; see below for details). Tissue was ground with two 4.5 mm steel balls unless otherwise stated, and all grinding steps consisted of 90 seconds of grinding at 850 RPM in a SPEX SamplePrep Geno Grinder 2010. Samples for the DNA extraction tests (Experiment I) were extracted according to the indicated protocol. All samples for PCR primer tests (Experiment II) were extracted using the Quick-DNA protocol. Details of each specific protocol follow.

#### Extract-N-Amp Plant PCR Kit(Sigma-Aldrich)

200 μl of Extraction Buffer buffer was added to the frozen samples, which were then ground as described above. Another 300 μl Extraction Buffer was then added (total 500 μl), mixed, and the solution incubated at 95° C for 10 minutes. 500 μl dilution buffer was then added before centrifuging the sample and transferring the supernatant to a clean tube.

#### PowerSoil DNA Isolation Kit (Molecular Biosciences; now Qiagen DNEasy PowerSoil Kit)

Soil samples were handled according to the manufacturer’s protocol, with two of the samples including 500 μl lysis buffer in the grinding step to test its effect. (It did not appear to alter extraction efficiency; data not shown.) Initial tests showed that the beads provided with the PowerSoil kit did not do a good job grinding plant leaf disks, so the grinding step of this protocol was modified as follows: 200 μl of lysis buffer from the PowerBead tubes was added to each sample tube (with two steel beads) and the tissue ground as above. 25 μl buffer C1 was added and mixed, and tubes centrifuged to pellet the debris. The supernatant was transferred to a clean tube and 100 μl buffer C2 was added. The remaining steps were carried out as per the manufacturer’s protocol, and DNA eluted in 100 μl.

#### DNeasy Plant kit (Qiagen)

100 μl (plants) or 200 μl (soils) of Buffer AP1 was added to the sample tubes prior to grinding as above. After grinding, additional Buffer AP1 was added to total 400 μl. For soil samples only, 4 μl RNase A was then added. The rest of the protocol followed manufacturer’s instructions.

#### Quick-DNA Fungal/Bacterial Miniprep Kit (Zymo Research)

200 μl BashingBead buffer was added to both plant and soil samples before grinding as described above. After grinding, 550 μl BashingBead buffer was added (total 750 μl) and the tube vortexed and centrifuged to pellet debris. The rest of the protocol was according to the manufacturer's instructions.

### Design of Maize-Specific Blocking Oligos

Blocking oligos for maize were designed following the procedure of (Agler et al. 2016). In brief, BLAST+ v2.2.31+ (Camacho et al. 2009) was used to align 30-bp sections of the maize mitochondria and chloroplast sequences to the Greengenes 97% OTU database v13.8 (DeSantis et al. 2006) (distributed as part of the qiime-default-reference v0.1.3 package for python; (Caporaso et al. 2010)). Sequences with the fewest Greengenes hits and a low probability of forming hairpins and homodimers (as determined by Primer3 (“Primer3” n.d.)) were chosen as new blocking oligos (Table S1, primers P10, P16, and P17).

### PCR Amplification

All samples had portions of the 16s gene amplified by PCR for sequencing. The exact protocol depended on the primer set being used. All primer and oligo sequences are in Supplemental Table 1. (Note that many of the primers include Illumina Nextera linkers used to prepare sequencing libraries.)

For the DNA extraction tests (Experiment I), soil samples were amplified with Universal primers (P01/P02) while the plant samples used the Discriminating primer set (P03/P04). PCR amplification tests (Experiment II) used the primer set marked for each sample, and each reaction was run in triplicate and then pooled for sequencing. A portion of each reaction was run on a 1% agarose gel to confirm amplification.

#### Universal primers

Universal primer reactions consisted of 1 μL of template DNA, 0.5 μL of forward primer P01 (515f; 0.2 μM final concentration), 0.5 μL of reverse primer P02 (806rB; 0.2 μM final concentration), 12.5 μL of 2x Hot Start Taq master mix (New England BioLabs), and 10.5 μL of water (=25 ul total volume). PCR amplification followed Earth Microbiome Project recommendations (“16S Illumina Amplicon Protocol: Earth Microbiome Project” n.d.): 94° C for 3 minutes; thirty cycles of 94° C for 45 s, 55° C for 60 s, and 72° C for 90 s; and a final 72° C extension for 10 minutes before holding at 4 C.

#### Discriminating primers

Discriminating primer reactions consisted of 1 μL of DNA template, 0.5 μL of forward primer P03 (799F; 0.2 μM final concentration), 0.5 μL of reverse primer P04 (1115R; 0.2 μM final concentration), 12.5 μL of 2x Hot Start Taq master mix (New England BioLabs) and 10.5 μL of water (= 25 μL total volume). The PCR program was 95° C for 5 minutes; thirty cycles of 95° C for 30 s, 55° C for 30 s, and 72° C for 30 s; and a final 72° C extension for 5 minutes before holding at 4 C.

#### PNA clamps

PNA clamps were stored at 100 μM at −20° C until use, during which time a small aliquot was diluted to 5 μM and stored at 4° C for up to one week. PCR reactions with PNAs were as the manufacturer’s recommendations, namely, 1 μL of template DNA, 1 μL of forward primer P01 (515f; 0.2 μM final concentration), 1 μL of reverse primer P02 (806rB; 0.2 μM final concentration), 1.25 μL mPNA (0.25 μM final concentration), 1.25 μL pPNA (0.25 μM final concentration), 12.5 μL of 2x Hot Start Taq master mix (New England BioLabs), and 7 μL of water (=25 μL total volume). The reaction mixture was placed in the thermocycler for PCR settings prescribed by PNA Bio: 94° C for 3 minutes; thirty cycles of 95° C for 15 s, 75° C for 10s, 55° C for 10 s, and 72° C for 60 s; and a final 72° C extension for 10 minutes before holding at 4° C.

#### Blocking Oligos

The blocking oligo reaction protocol was adapted from (Agler et al. 2016) and consists of two PCR steps. (Primer names are given at the end because they varied by species and target region, but the methodology was the same.) First-step amplification consisted of 0.5 μL of template DNA, 0.16 μL of forward primer (0.08 μM final concentration), 0.16 μL of reverse primer (0.08 μM final concentration), 0.5 μL of forward blocking oligo (0.25 μM final concentration), 0.5 μL of reverse blocking oligo (0.25 μM final concentration), 10 μL of 2x Hot Start Taq master mix (New England BioLabs) and 8.18 μL of water (= 20 μL total). The first amplification reaction mixture was placed in the thermocycler for 95° C for 40 s; ten cycles of 95° C for 35 s, 55° C for 45 s, and 72° C for 15 s; and a final 72° C extension for 2 minutes before holding at 4° C.

First-round PCR product was subjected to an enzymatic cleanup by adding 1μL Antarctic phosphatase (New England Biolabs), 1μL Exonuclease I (New England Biolabs), and 2.44μL 10x Antarctic phosphatase buffer. The reaction was incubated at 37° C for 30 minutes followed by 85° C for 15 minutes to inactivate the enzymes. Reactions were centrifuged at 7000 rpm until a white pellet formed on the wall of the tube, and 10μL of the supernatant was transferred to a fresh PCR tube.

Second-step PCR amplification consisted 2.0 μL of DNA from step 1, 0.83 μL of forward primer (0.166 μM final concentration), 0.83 μL of reverse primer (0.166 μM final concentration), 25 μL of 2x Hot Start Taq master mix (New England BioLabs) and 21.34 μL of water (=50 μL total volume), with the amplification program of 95° C for 40 s; twenty-five cycles of 95° C for 35 s, 55° C for 45 s, and 72° C for 15 s; and a final 72° C extension for 2 minutes before holding at 4° C. Each reaction was run in triplicate and then pooled for sequencing.

PCR primer varied by species and target region. For all samples, BO_3/4 used amplification primers P05 (B341F) and P06 (B806R) in both PCR steps. The soybean samples used the default chloroplast-blocking oligos from (Agler et al. 2016) (P07 [3C30-F]) & P08 [c11BV3-R]) in the first PCR, while all other samples used a maize-specific forward blocking oligo (P09) with the default reverse (P08 [c11BV3-R]). For technical reasons, some of the BO_5/7 reactions used amplification primers with linkers in the first PCR (P10 [799F] & P11 [1192R]), while others used ones without linkers (P12 [799F] & P13 [1192R]); all used primers P10 and P11 in the second step. This information is recorded in the sample keys (part of the analysis pipeline at https://www.github.com/wallacelab/paper-giangacomo-16s-methods) and did not cause any significant differences in the results (data not shown). (Also note that even though both are called 799F in the literature, primers P10 and P12 have a slightly different sequence from primer P03.) For BO_5/7, mitochondria-blocking oligos for soybean in the first amplification step were from (Agler et al. 2016) (P14 [5M30-F] & P15 [cl1BV5-R]), while all other samples used maize-specific ones (P16 & P17).

### Library preparation

For the DNA extraction tests (Experiment I), PCR products were purified with a QIAQuick PCR Purification kit (QIagen) according to the manufacturer’s instructions; those for PCR amplification tests (Experiment II) were purified with AGENCOURT AMPure XP magnetic beads. Illumina sample-specific barcodes and sequencing adapters were added in a final PCR reaction consisting of (per sample) 10 μl 2x Hot Start Taq master mix (New England BioLabs), 2 μl each of Nextera N7 and S5 barcoding primers (Illumina), 1 μl purified template, and 5 μl water (20 μl total). The PCR reaction cycle was 95° C for 5 minutes; eight cycles of 95° C for 30 s, 55° C for 30 s, and 72° C for 30 s; and a final 72° C extension for 5 minutes before holding at 4° C.

Barcoded samples were sent to the Georgia Genomics and Bioinformatics Core at the University of Georgia for quantification, normalization, pooling, and sequencing on an Illumina MiSeq. Paired-end 2×250 sequencing was performed for DNA extraction tests (Experiment I). Since the Blocking Oligos protocol resulted in longer amplicons, paired-end 2×300 was used for the PCR amplification tests (Experiment II). All read data are available at the NCBI Sequence Read Archive, Bioproject PRJNA646931.

### Bioinformatic processing

Raw sequencing reads were processed down to amplicon sequence variants (ASVs) using the QIIME 2 pipeline (version 2019-7; (Bolyen et al. 2019)). DNA extraction tests were processed separately from PCR amplification tests, and different primer sets were processed separately within each test set, though all followed a common pipeline. All bioinformatic scripts are available at https://www.github.com/wallacelab/paper-giangacomo-16s-methods.

First the final 100 (extraction tests) or 150 (primer tests) base pairs of reverse reads were removed with cutadapt (command ‘cutadapt --cut’; (Martin 2011)) because initial checks found that the high error rate in these regions interfered with read joining. Primers were removed with cutadapt (command ‘qiime cutadapt trim-paired’, with forward and reverse primer sequences supplied), followed by joining with vsearch (‘qiime vsearch join-pairs’; (Rognes et al. 2016)). Joined pairs were quality filtered with ‘qiime quality-filter q-score-joined’.

Amplicon sequence variants were called with Deblur (‘qiime deblue denoise-16s’; (Amir et al. 2017)), with trim lengths based on the primer set used (250 bp for Universal and PNAs, 295 for Discriminating, 370 for Blocking Oligos V5-V7, and 400 for BO V3-V4) and keeping all ASVs with at least 2 reads across any samples (‘--p-min-reads 2 --p-min-size 1’). Taxonomy was assigned by extracting the targeted regions from the Silva 132 0.99 OTU database (Quast et al. 2013) (‘qiime feature-classifier extract-reads’), training a custom naive bayes classifier (‘qiime feature-classifier fit-classifiers-naive-bayes’), and using this classifier to assign taxonomy to the ASVs (‘qiime feature-classifier classify-sklearn’; (Pedregosa et al. 2011; Bokulich et al. 2018)).

Since different primer sets targeted different parts of the 16s gene, a phylogenetic tree containing all samples in a test set could not be made directly from the ASVs. Instead, we used BLAST+ v2.2.31 (Camacho et al. 2009) to identify the best hit for each ASV in the Silva 132 0.99 OTU database (Quast et al. 2013), and that hit was used to anchor each ASV in the corresponding Silva phylogenetic tree using the R package ape v5.3 (Paradis and Schliep 2019).

### Data unification and filtering

Treatment comparisons were performed by exporting the Qiime2 artifacts to standard file formats (BIOM (McDonald et al. 2012) for ASV tables, FASTA for sequences, etc.). Phyloseq v1.28.0 (McMurdie and Holmes 2013) was then used to combine ASVs from each primer set into a single unified data table containing ASV counts, sample metadata, assigned taxonomy, and phylogeny, which was then collapsed to the Genus level to allow comparison across primer sets. (This reduced ASVs to the “operational phylogenetic units,” or OPUs, of (Agler et al. 2016)). Samples were then filtered to with a minimum depth of 1000 (Experiment I) or 500 (Experiment II), at least 3 reads per OPU, and each OPU present in at least 2 samples.

### Statistical analyses

<u>Organellar contamination</u> of primer sets was tabulated by summing the read counts in each sample with the taxonomic assignment of Chloroplast (order) or Mitochondria (family). Organelles were then removed from the data before performing other analyses.

<u>Alpha Diversity</u> of DNA extraction tests was calculated in phyloseq using the plot_richness() function for observed OTUs and Shannon diversity.

<u>Shared and unique OPUs</u> were calculated by converting reads counts to presence/absence data. We then tabulated the OPUs unique to a given sample set or shared with at least one other sample set.

<u>Principal coordinate plots</u> were generated by rarefying samples to 2000 (Experiment I) or 500 (Experiment II) reads and calculating the weighted unifrac distance with rbiom 1.0.0 (Smith 2019). (Phyloseq v1.28.0 appears to have a bug in its unifrac calculation; see https://github.com/joey711/phyloseq/issues/936). The cmdscale() function in R was then used to convert the distance matrices to principal coordinates.

<u>Taxonomic distortion</u> was calculated by using the phyloseq tax_glom() function to collapse the data at various taxonomic levels (phylum, class, etc.) and the phyloseq_to_deseq2() function to convert it to a DESeq2 (Love, Huber, and Anders 2014) object with both sample type and treatment as design variables. Since DESeq2 ignores any taxa with 0 reads in any samples, if any collapsed taxon had 0 counts, a pseudocount of 1 was then added to all counts. Data was rarefied to the lowest read count among samples, and the DESeq() function was used to find significantly different taxa. (Rarefaction is usually not recommended for DESeq data, but we found that without it the program tended to find most taxa in a sample changing in a single direction that correlated with the total read depth of that sample.) DESeq significance was determined with a Wald test statistic and parametric fit for all levels except Domain, which was fit with a mean-dispersion estimate. (This difference was due to a consistent error when trying to fit Domain with a parametric dispersion estimate, possibly due to there being only 2 domains [Bacteria/Archaea] and Archaea having no reads in many samples.) Taxa were considered significantly distorted if their adjusted p-value was below 0.01.

<u>Verification of sample identity</u> was performed using both BLAST (Camacho et al. 2009) and Kraken2 (Wood, Lu, and Langmead 2019). BLAST analysis was done by taking the first 1000 reads of each raw fastq file, BLASTing them against the Silva 132 0.99 OTU database (Quast et al. 2013), and tallying up the species of all hits corresponding to ‘Chloroplast’ or ‘Mitochondria’. Kraken2 analysis involved using Kraken2 to assign all reads in each sample to a taxon based on a custom database consisting of the NCBI plants database. (Reads that did not match the plant database--such as bacterial reads--were simply “unclassified.”)

<u>Plots</u> were generated with ggplot2 (Wickham 2016) and minor cosmetic adjustments made in Inkscape (Developers n.d.).

### Software packages

The following software packages were used in this analysis:

- R packages ape v5.3 (Paradis and Schliep 2019), argparse v2.0.1 (Davis 2019), DESeq2 v1.24.0 (Love, Huber, and Anders 2014), dplyr v0.8.3 (Wickham et al. 2019), ggplot2 v3.2.1(Wickham 2016), gridExtra v2.3 (Auguie 2017), igraph v1.2.5 (Csardi, Nepusz, and Others 2006), phyloseq v1.28.0 (McMurdie and Holmes 2013), rbiom v1.0.0 (Smith 2019), tidyr v1.0.0 (Wickham and Henry 2019), vegan v2.5.5 (Oksanen et al. 2019)
- Python packages argparse v1.1, biopython v1.74 (Cock et al. 2009), matplotlib 3.1.0 (Hunter 2007), primer3-py v0.6.0 (“primer3-Py” n.d.)
- Command-line tools biom v2.1.7 (McDonald et al. 2012), blast+ v2.2.31 (Camacho et al. 2009), Conda v4.8.2, Clustal Omega v1.2.1 (Sievers et al. 2011), cutadapt v2.4 (Martin 2011), libprimer3 v2.5.0 (“Primer3” n.d.), QIIME2 v2019-7 (Bolyen et al. 2019), vsearch v2.7.0 (Rognes et al. 2016)

## Results

### Experiment I - DNA Extraction

We tested four different commercial extraction methods covering a range of manufacturers and target tissues (soil, microbial, plant) to determine how well each captures the microbial community. Three of these--MoBio Powersoil, Qiagen DNeasy Plant, and Zymo Quick DNA-- consisted of column-based affinity purification, and one phase-separation method (Extract-N-Amp) was included for comparison. Since the true composition of the community is unknown, the MoBio (now Qiagen) PowerSoil kit recommended by the Earth Microbiome Project was used as the baseline. (The recommendation was later updated to the magnetic bead-based version of this kit; (Marotz et al. 2017)). Samples consisted of leaves from corn, soybean, and Arabidopsis, and also a soil sample. Since the results of the primer tests (Experiment II) were not known at the time, plant samples were amplified with chloroplast-discriminating primers and the soil samples were amplified with Earth Microbiome Project universal primers (see Methods).

Since the samples were the same across all extraction methods, we expect that the “best” methods will recover more taxa (=higher alpha diversity). However, no single method followed this pattern for all sample types (Figure 1). The Power Soil and EasyDNA kits recovered the highest alpha diversity of soil samples, as might be expected since the other two kits had not been designed with soil in mind. However, in plant samples we found that the DNeasy Plant kit performed best for Arabidopsis, EasyDNA worked best for maize, and everything performed roughly evenly for soybean. The number of unique taxa found with each method (Supplemental Figure S1) reinforces this, as the best method for a sample usually has the largest number of unique taxa, while others share most of their taxa with at least one other method.

**Figure 1.**
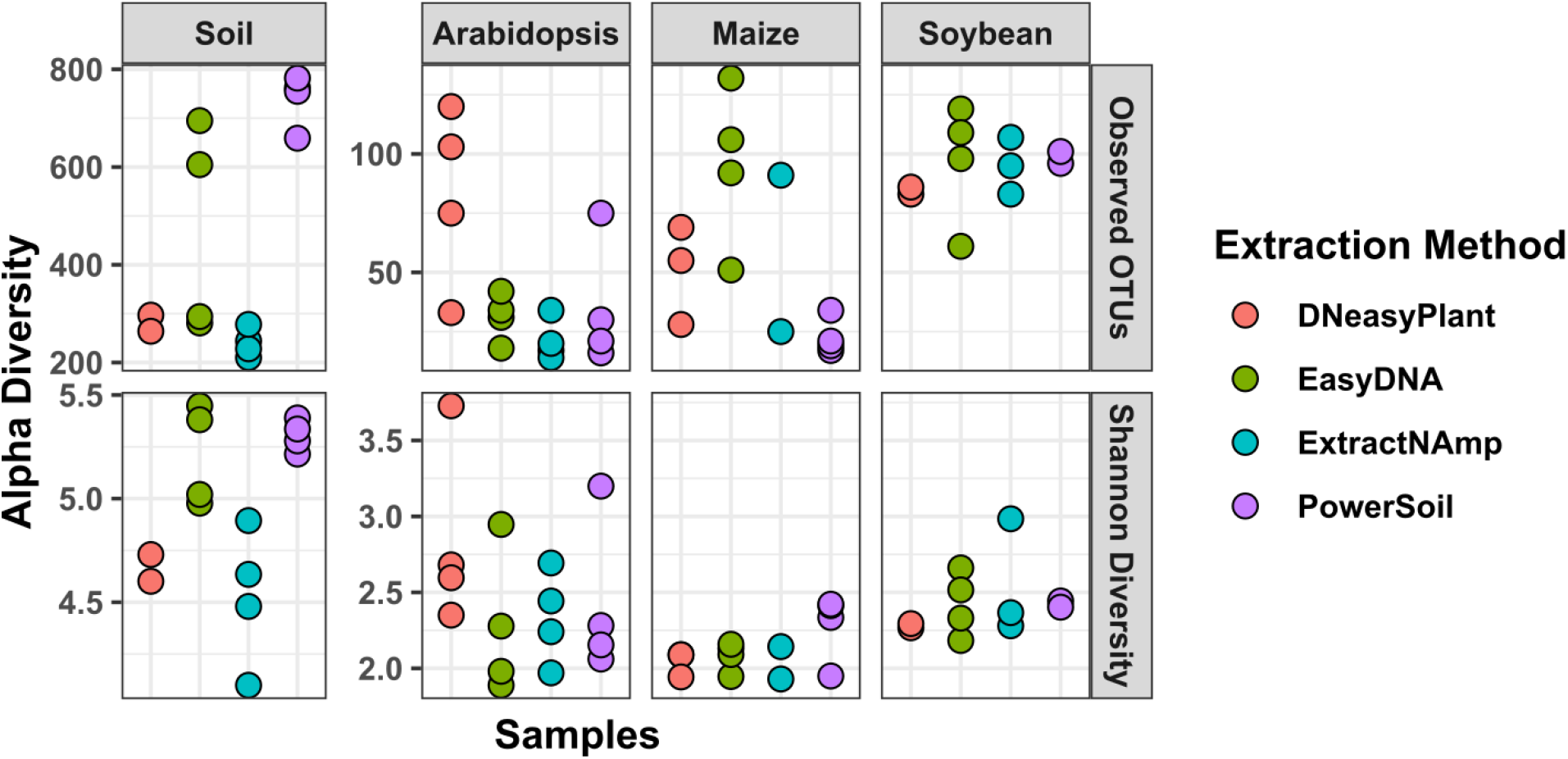
Alpha diversity of extraction methods. The total number of observed OTUs and the Shannon diversity of each extraction method on different samples is shown; each point represents one sample. Different kits appear better at capturing larger diversity depending on the sample type (soil vs. leaf) or even the plant species involved.

Overall community composition was assessed with weighted UniFrac analysis. As expected, samples generally cluster by sample type (Figure 2a), except that the maize samples extracted with PowerSoil cluster with the Arabidopsis samples. Analysis of the raw data indicates that this was not due to a labeling error (Supplemental Figure S2), so it appears that the PowerSoil kit has a strong effect on the recovered maize microbiome but not on soybean and Arabidopsis. Within each sample type, soil shows the most distinct clustering by extraction method (Figure 2b-e). The three plant samples show less distinct clustering of all methods from each other, although maize leaves showed a very strong separation of PowerSoil extractions from the other three, as mentioned earlier. If we take the PowerSoil kit to be the current standard, none of the other methods consistently overlapped with it, although the EasyDNA kit came closest in all samples but maize. (And in that case the PowerSoil kit is likely the one with stronger biases.)

**Figure 2.**
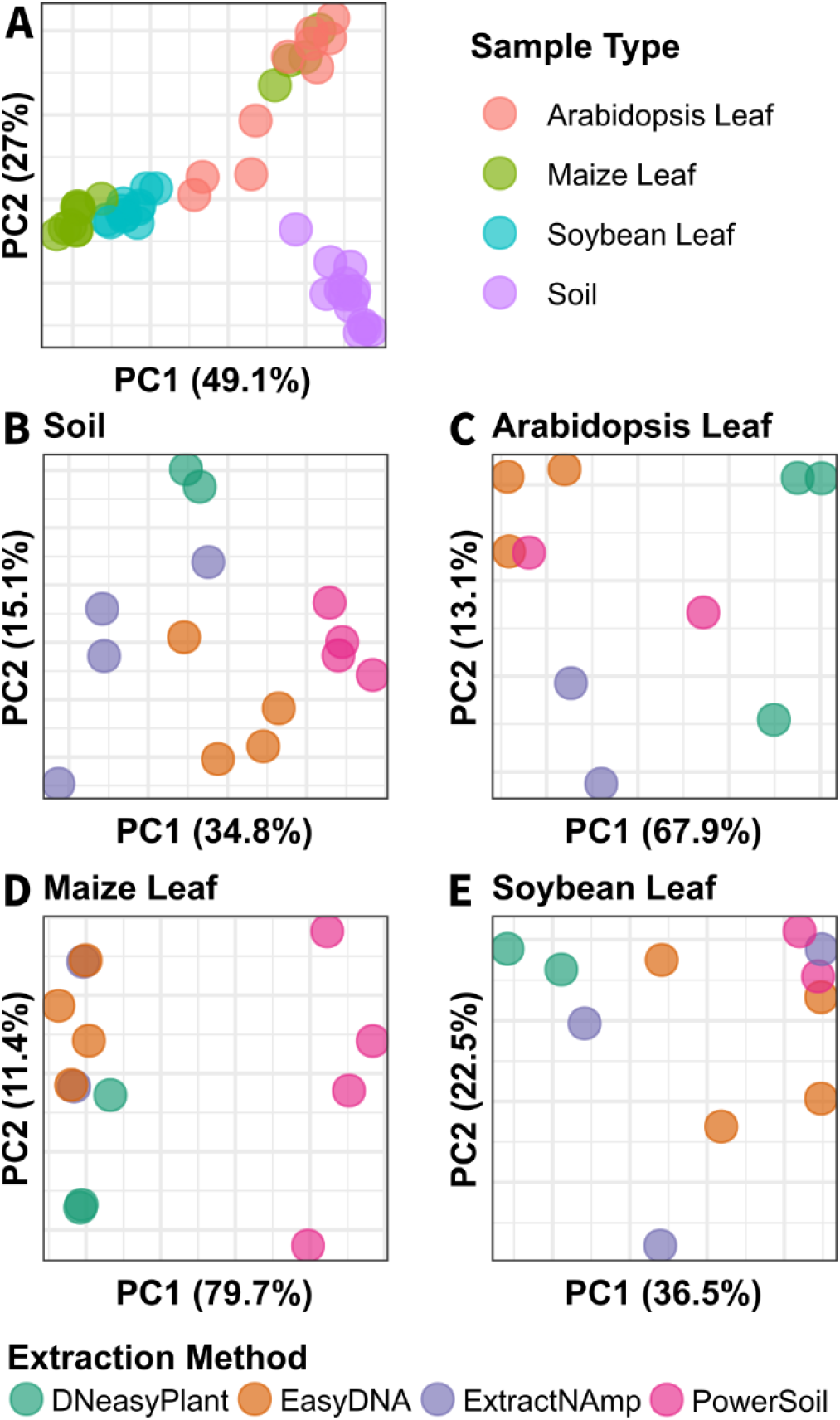
Principal coordinates of extraction methods. The principal coordinates of of all samples (A) or individual sample types (B-E) were calculated using the weighted Unifrac distance metric (Lozupone and Knight 2005). The key for part (A) is above, while the key for the other four panels is below. None of the extraction methods consistently overlaps with the others, indicating that each method shows a slightly different version of the underlying community.

To determine just how different methods compared at recovering different bacterial taxa, we used DESeq2 (Love, Huber, and Anders 2014) to determine which groups at various taxonomic levels were distorted relative to the Power Soil method (the current community standard) (Figure 3). This clearly shows that the EasyDNA method has the least distortion relative to PowerSoil, with most distortion limited to a single genus (*Aureimonas*) within the Proteobacteria. The DNEasy Plant kit showed significant distortion, with the enrichment of several Proteobacteria clades (especially the genus *Sphingomonas*) and depletion of Actinobacteria (genera *Cutibacterium* and *Corynebacterium*) and the Firmicutes family Staphylococcaceae. A complete table of all significantly distorted clades is in Table S2.

**Figure 3.**
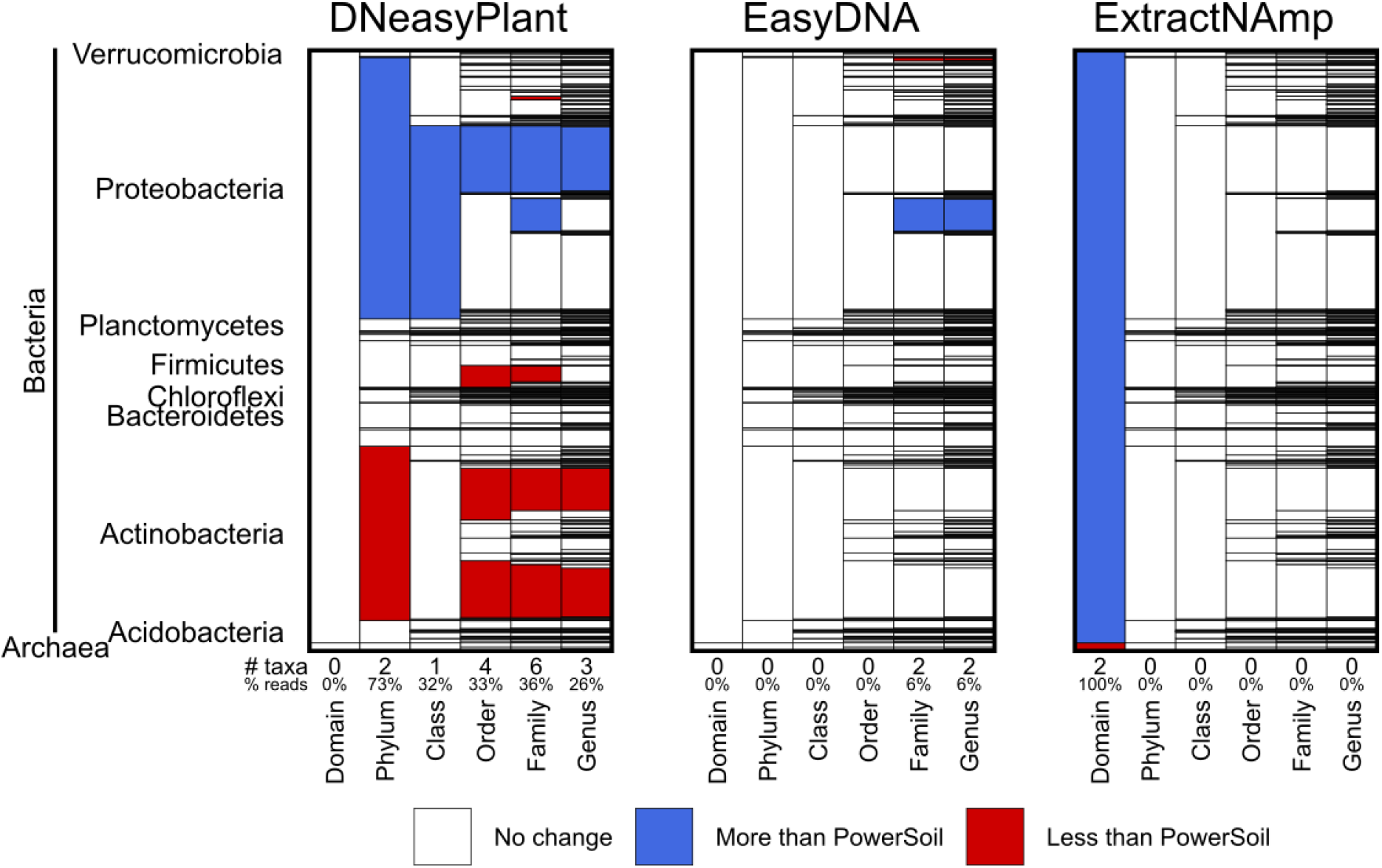
Distortion of the community relative to PowerSoil extraction. Each plot shows stacked rectangles corresponding to bacterial & archaeal taxa going from general (Domain, left) to specific (Genus, right), with bar size proportional to the number of overall reads from that taxon across all samples. DESeq2 (Love, Huber, and Anders 2014) was used to test for significant distortion at each taxonomic level, and significantly distorted taxa (adjusted p-value <= 0.01) are colored, with blue indicating more relative counts and red indicating taxa with fewer relative counts. The names of major Bacterial phyla are to the left, and the number of distorted taxa and percent of reads in affected taxa is shown at bottom.

### Experiment II - Amplification

We compared 3 different methods to exclude organellar DNA from 16s amplification of plant samples: discriminating primers, PNA clamps, and blocking oligos targeting two different regions of the 16s rRNA gene (V3-V4 and V5-V7; see Methods). Each was compared with generic universal primers from the Earth Microbiome Project as the control. Samples tested were two soil samples, a defined 20-member community (ATCC 20-Strain Even Mix Genomic Material), and two plant leaf samples (maize and soybean). The soil and defined community samples were used to detect bias in the primer sets, as they should not contain organelles.

As expected, the universal primers amplified almost pure organellar DNA from plant samples (Figure 4). Discriminating primers and the blocking oligos targeting the V5-V7 region (BO_5/7) had the lowest organellar sequence (nearly zero), while the PNA clamps and V3-V4 blocking oligos (BO_3/4) had intermediate levels.

**Figure 4.**
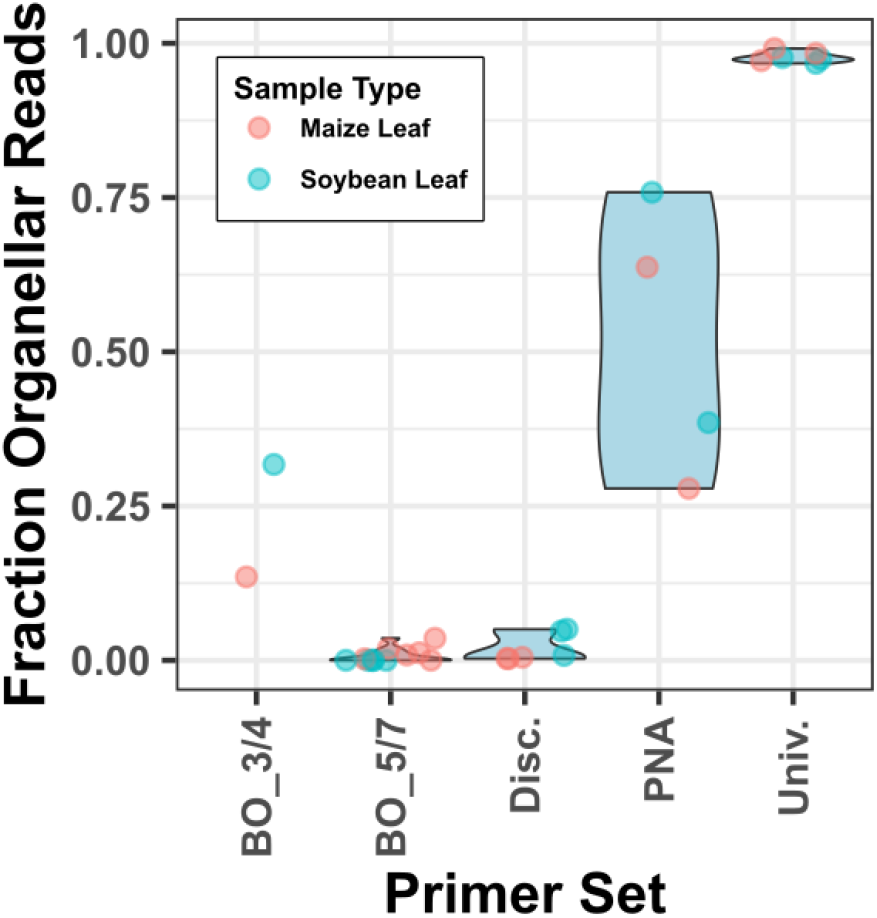
Organelle contamination by primer set. The fraction of recovered reads that match chloroplast or mitochondrial 16s sequences is shown for the 5 amplification method sets when used on maize or soybean leaf tissue. Blocking oligos BO_5/7 and discriminating primers have low organellar contamination, BO_3/4 and PNAs intermediate, and Universal primers high. BO, blocking oligos; Disc., discriminating; Univ, universal.

To determine how much each method distorts the view of the underlying community, we used them to amplify bacterial communities from two soil samples and a defined microbial community (Figure 5). In all three samples, the PNAs and Universal primer set cluster together (though not quite on top of each other with soil). Interestingly, the V5-V7 blocking oligos cluster with the Universal primers when using a defined community, as reported in the original publication, but they are strongly distinct in soil samples, indicating that there is still significant distortion occurring. The discriminating primers and V3-V4 Blocking Oligos always cluster separately, indicating significant distortion.

**Figure 5.**
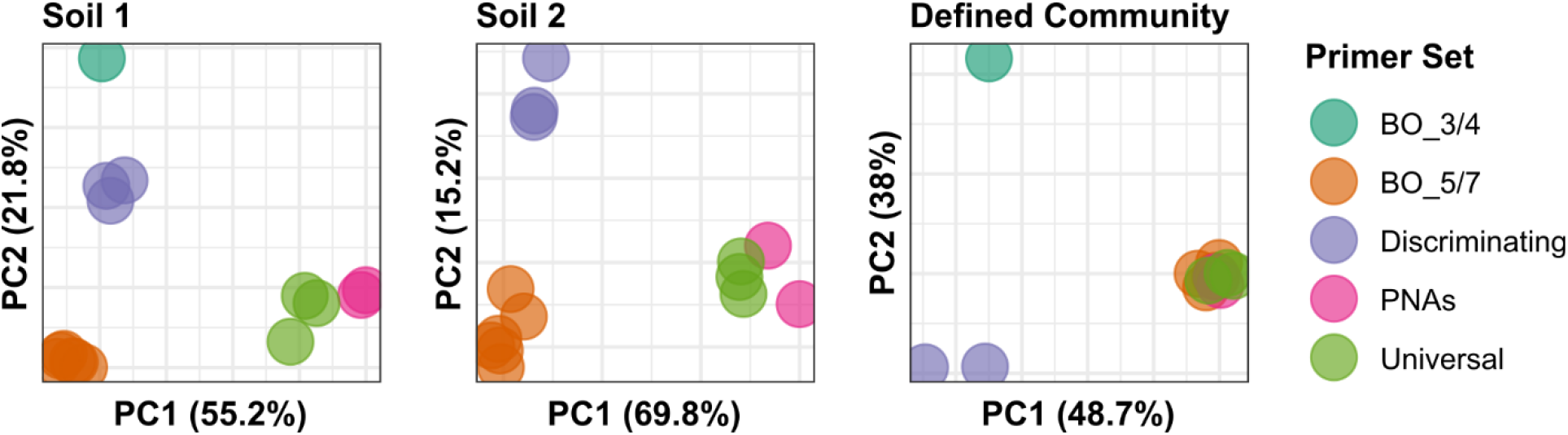
Principal Coordinates of primer sets. The weighted UniFrac distances of both soil samples and a 20-member defined community are shown. PNAs generally cluster close to (but not quite overlapping) the Universal primers, but the other three primer sets are generally separate. Interestingly, the Blocking oligos 5/7 set appears to exactly overlap PNAs and the Universal primers in a defined community (matching a claim from the original publication (Agler et al 2016)), but they are very different with the more complex soil samples.

To identify specific bacterial clades that are affected by the different primer sets, we determined the significantly different taxa with DESeq2 (Love, Huber, and Anders 2014). The Universal primer was set as the reference, and we limited the analysis to just the soil samples (since the Universal set has almost no bacterial reads on leaf samples) (Figure 6). As expected, PNAs show no distortion relative to the Universal primer set. The other three sets show large numbers of taxa being distorted, including the Archaea and most clades within the Bacteria. All distorted bacterial clades are too numerous to mention (see Supplemental Table S3 for the full list), though in broad strokes Proteobacteria are enriched; Verrucomicrobia, Chloroflexi, and Acidobacteria are depleted; and clades within the Actinobacteria can go either way. The distortion pattern is similar between the BO_5/7 and Discriminating sets, probably because they share a primer site (799F), though it is surprising that the BO_3/4 set also shows a broadly similar pattern of distortion. A full list of distorted taxa is in Supplemental Table S3.

**Figure 6.**
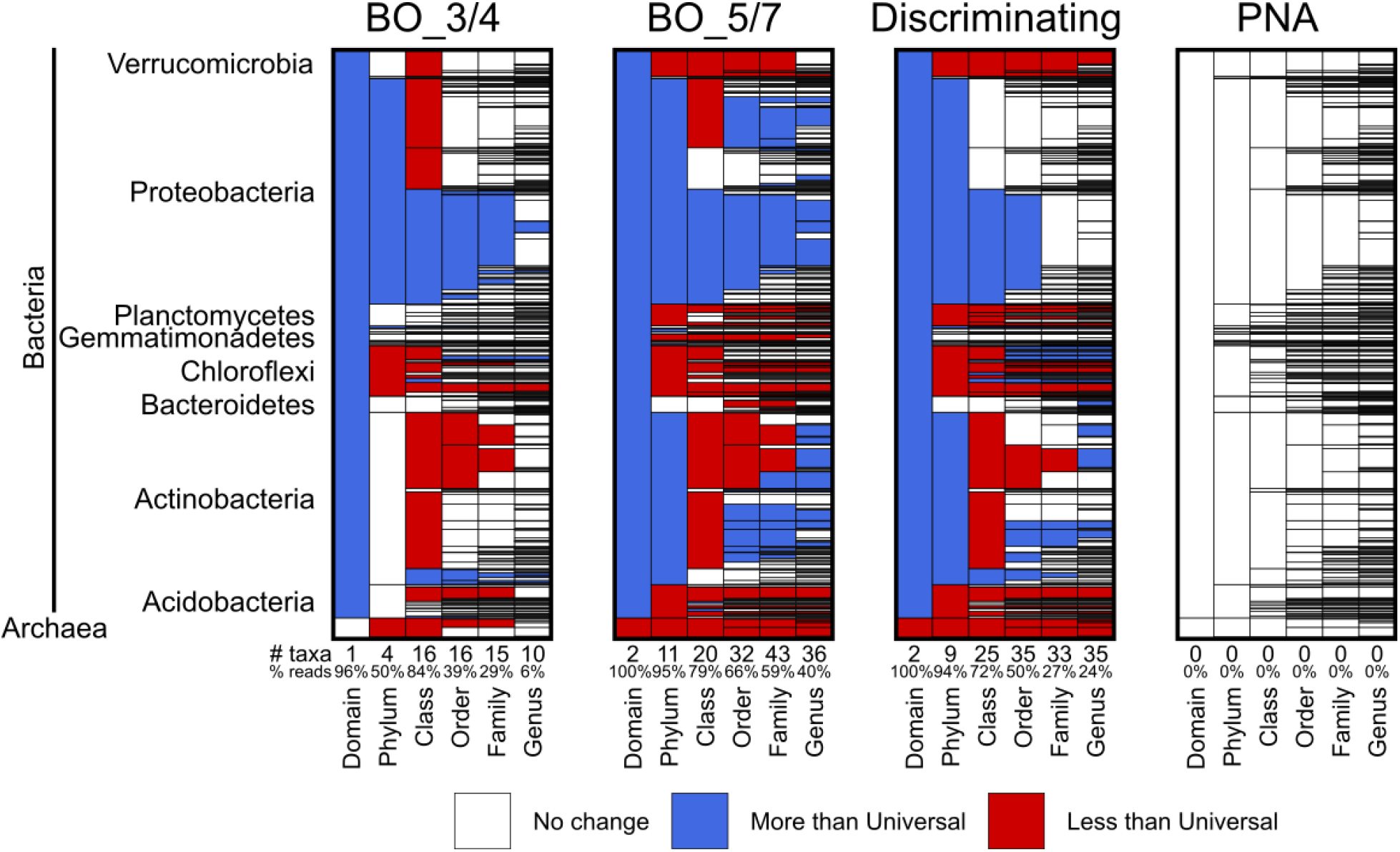
Distorted taxa of amplification protocols relative to the Universal primer set. Plot layout is the same as per Figure 3. PNAs show no significant distortion relative to the Universal primer set, while all three other methods show significant distortion across many taxa.

When looking at the total number of genera recovered by each method (Supplemental Figure S3) only the V3-V4 Blocking Oligos show significantly lower genus counts with the defined community. In soil samples, however, the PNA and Universal primer sets recover dramatically more genera than the other methods. On leaves, the Discriminating Primers and PNA recover the most genera; the low number of genera for the Universal primer set is likely due to almost all of its reads coming from chloroplasts instead of bacteria.

## Discussion

All current microbiome methods are known to result in some degree of bias (Pollock et al. 2018). Since removing all bias is impossible, current best practices are to treat everything within an experiment as identically as possible so that all samples at least suffer the same biases. Nonetheless, researchers generally want to reduce bias as much as possible to get the clearest results they can. With that in mind, our results suggest the following recommendations regarding DNA extraction and amplification of plant-associated samples.

### Choice of extraction method (Experiment I)

The extraction methods we tested were chosen based on their prevalence in the field and/or ease of use. Principal coordinate analysis shows a distinct clustering for each method, confirming that each has unique biases (Figure 3). For soil samples (and by extension, rhizosphere samples), the best extraction methods were unsurprisingly those designed for soil extraction: the PowerSoil and EasyDNA kits. These kits had the highest recovered alpha diversity from soil (Figure 1), the largest number of recovered OTUs, most of which were shared between them (Supplemental Figure S1), were relatively close to each other in overall community composition (Figure 2), and showed relatively minor taxonomic distortion (Figure 3).

The choice of extraction method for plant samples is more complex. For Arabidopsis, the DNeasy Plant kit had the greatest alpha diversity and number of recovered genera, while for maize it was the EasyDNA kit, and soybean showed no clear winner. A challenge with plant samples is that the extraction method must not only lyse the microorganisms (including endophytes within the host tissue), it must also remove the phenolic compounds and other inhibitors of downstream processing steps (Pollock et al. 2018). These compounds vary from species to species, so it may be that some testing is required to identify the best method for each species. Without that data--or for studies that will span a wide range of species--the best advice is again to choose a single method and use it consistently, but be aware of host-induced biases that can occur. As for which to choose, from this (admittedly small) dataset it appears that the EasyDNA may be the most middle-of-the-road choice, avoiding both the exceptional good result of the DNEasy Plant kit on Arabidopsis (Figure 1) and the outlier result of PowerSoil on maize (Figure 3).

When dealing with multiple sample types such as bulk soil, rhizosphere, stem, and leaf, the choice of method should be guided by the sample and goals of the experiment. Any conditions that will be compared should be extracted with the same method. This is necessary to avoid condition-specific biases when comparing, for example, bulk soil to rhizosphere or either of those to stem tissue to trace the origin of plant endophytes. If, however, there is no reason to compare across compartments, then each compartment can be extracted with the method that works best for it. This will recover a better snapshot of the community in each compartment, but at the cost of not being able to compare between them. It also adds more complexity to the experiment that could result in an error, such as by using the wrong method on some of the samples.

### Choice of 16s amplification method (Experiment II)

The goal of choosing an amplification protocol is to minimize bias while also minimizing the amount of host DNA in the sample. Host contamination is not generally a problem for soil samples, but becomes crucial for samples composed mostly of host tissue (roots, stems, leaves, etc.).

Of the methods we tested, the Blocking Oligos and Discriminating Primers did the best job at excluding DNA from organelles, excluding almost all organellar DNA (Figure 4). BO_3/4 was more modest, at ~25% organellar reads. The PNA clamps also showed only modest control (30-75% organelle DNA), though it should be noted that we tested only a single concentration of PNAs; higher concentrations might be able to exclude more plastid sequence, but at the cost of using more PNA reagent per reaction.

Comparing amplification of non-plant samples (two soils and a defined community) revealed that only the PNAs maintained a community composition comparable to the Universal controls (Figure 5). The PNA method was also consistently second-best in terms of number of bacterial genera recovered (Supplemental Figure S3), coming behind the Universal primers for soil samples and the Discriminating primers for plant samples. Both sets of Blocking Oligos performed consistently poorer than the other methods. We also found them much more difficult to work with, with a more complex protocol and a much higher failure rate than the other methods (personal observation).

Given these results, we recommend to use either PNA clamps or plastid-discriminating primers. The PNA clamps are preferred because they minimally distort the underlying community while still increasing target sequences ~20-fold; higher concentrations of PNA may be able to improve this further (but at increased reagent cost). Discriminating primers are slightly easier and less expensive, but do result in significant taxonomic distortion. We do not recommend blocking oligos for most labs because they are very species-specific, have a more complex protocol, and appear to distort the taxonomic profile (Figure 5). The V3-V4 set used here and recommended in the original paper also requires longer read lengths than most other methods (paired-end 300 instead of paired-end 250), which also increases costs and limits which Illumina sequencing platforms can be used. Blocking oligos might still be useful for laboratories that focus on a single species (like *Arabidopsis*) and that are willing to invest the resources to optimize and confirm the procedures; otherwise we recommend a simpler protocol.

### Implications for plant-associated microbiome work

Plant-associated microbiome analysis presents unique challenges due to the abundance of host-associated DNA, especially from plant organelles. Probably for this reason, the first release of the Earth Microbiome Project data included almost no plant-associated samples (Thompson et al. 2017). However, plant-associated microbes play crucial roles in global ecosystems and agriculture. Understanding these interactions will be necessary to fully harness them during the 21st Century. Although several of the methods compared in this paper help probe the plant microbiome, better methods are still needed. Critically, to our knowledge there are no current methods that reliably separate plant host DNA from microbes, a necessary step for whole-metagenome profiling of the plant microbiome (especially the plant endosphere). We urge the development of these methods, and expect that their creation will help unveil unexpected aspects of plant-microbe interaction.

## Acknowledgements

We thank Dr. Matthew Agler for help troubleshooting the Blocking Oligos protocols. This work was supported by startup funding from the University of Georgia.

