## Supplemental data and files for "Comparing DNA Extraction and 16s Amplification Methods for Plant-Associated Bacterial Communities": 16s methods paper - Supplemental Data v1.0.pdf

**Supplemental Table S1 – Primers and oligos used in this study**

| Primer set | Name | ID* | Sequence† | Reference |
|---|---|---|---|---|
| Universal, PNAs | 515f | P01 | [TCGTCGGCAGCGTCAGATGTGTATAAGAGACAG]GTGYCAGCMGCCGCGGTAA | (16S Illumina Amplicon Protocol : Eart...) |
| Universal, PNAs | 806rB | P02 | [GTCTCGTGGGCTCGGAGATGTGTATAAGAGACAG]GGACTACNVGGGTWTCTAAT | (16S Illumina Amplicon Protocol : Eart...) |
| Discriminating | 799F | P03 | [TCGTCGGCAGCGTCAGATGTGTATAAGAGACAG]ACCMGGATTAGATACCCKG | (Shade et al. 2013) |
| Discriminating | 1115R | P04 | [GTCTCGTGGGCTCGGAGATGTGTATAAGAGACAG]AGGGTTGCGCTCGTTG | (Shade et al. 2013) |
| PNAs | pPNA | pPNA | GGCTCAACCCTGGACAG‡ | (Lundberg et al. 2013; PNA PCR Blockers \| PNA BIO ) |
| PNAs | mPNA | mPNA | GGCAAGTGTTCTTCGGA‡ | (Lundberg et al. 2013; PNA PCR Blockers \| PNA BIO ) |
| Blocking Oligos | B341F | P05 | [TCGTCGGCAGCGTCAGATGTGTATAAGAGACAG]CCTACGGGAGGCAGCAG | (Agler et al. 2016) |
| Blocking Oligos | B806R | P06 | [GTCTCGTGGGCTCGGAGATGTGTATAAGAGACAG]GGACTACHVGGGTWTCTAAT | (Agler et al. 2016) |
| Blocking Oligos | 3C30-F | P07 | GAGGTAGAAGGCCTACGGGTCCTGAACTTC | (Agler et al. 2016) |
| Blocking Oligos | cl1BV3-R | P08 | TGTCAGTGTCGGCCCAGCAGAGTGCTTTCG | (Agler et al. 2016) |
| Blocking Oligos | Maize chloroplast blocking oligo Fwd | P09 | GAGGTGGAAGGCCTACGGGTCGTCAACTTC | This study |
| Blocking Oligos | 799F (with linker) | P10 | [TCGTCGGCAGCGTCAGATGTGTATAAGAGACAG]AACMGGATTAGATACCCKG | (Agler et al. 2016) |
| Blocking Oligos | 1192R (with linker) | P11 | [GTCTCGTGGGCTCGGAGATGTGTATAAGAGACAG]ACGTCATCCCCACCTTCC | (Agler et al. 2016) |
| Blocking Oligos | 799F | P12 | AACMGGATTAGATACCCKG | (Agler et al. 2016) |
| Blocking Oligos | 1192R | P13 | ACGTCATCCCCACCTTCC | (Agler et al. 2016) |
| Blocking Oligos | 5M30-F | P14 | AGATCAGGGGCTCAGCTAACGCGTGAACAC | (Agler et al. 2016) |
| Blocking Oligos | cl1BV5-R | P15 | TTTTGGCAGGGCGTACTAAACCCACTTACT | (Agler et al. 2016) |

| | | | | |
|---|---|---|---|---|
| Blocking Oligos | Maize mitochondria blocking oligo Fwd | P16 | ACTAGGTGCTGTGCGACTCGACCCGTGCAG | This study |
| Blocking Oligos | Maize mitochondria blocking oligo Rev | P17 | TGTTCAGGGTTCCAAACTCATAGTGGCAAC | This study |

** The Primer ID used in the Methods section*

*† Sequence in brackets indicates Illumina-compatible linkers to add sample barcodes in the final library prep PCR.*

*‡ These oligos are PNAs (Peptide Nucleic Acids), not standard DNA*

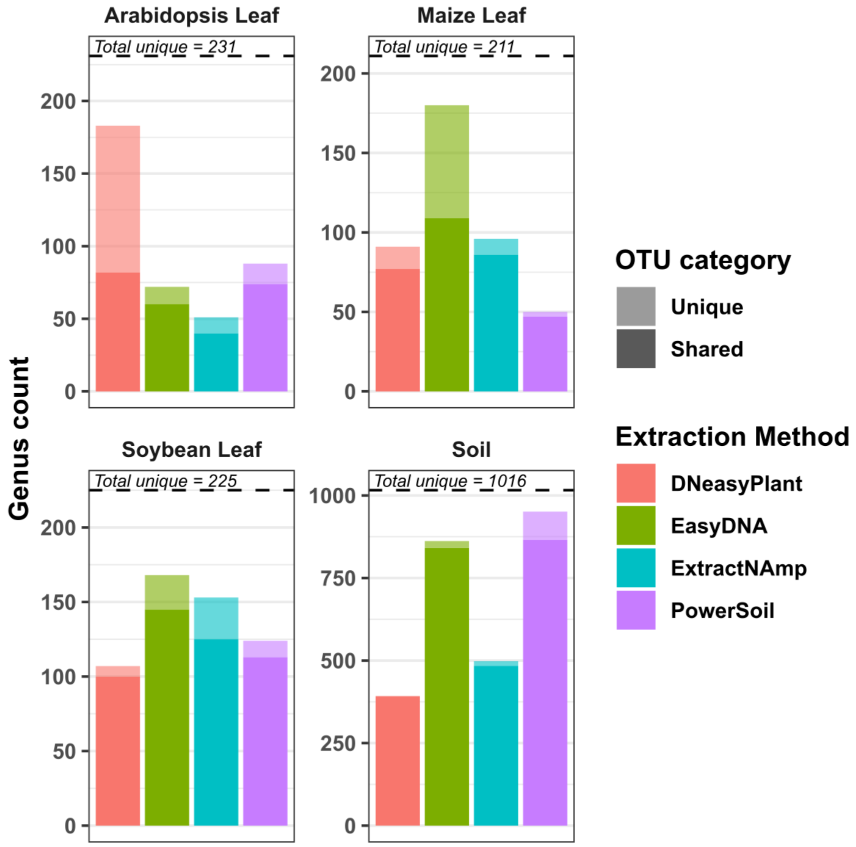


**Supplemental Figure S1 – Shared and unique genera among extraction methods.** The number of shared (dark) an unique (light) genera identified by each method is shown relative to the total number of unique genera (dotted line). A genus is counted as "shared" if at least two methods identified members of that genus.

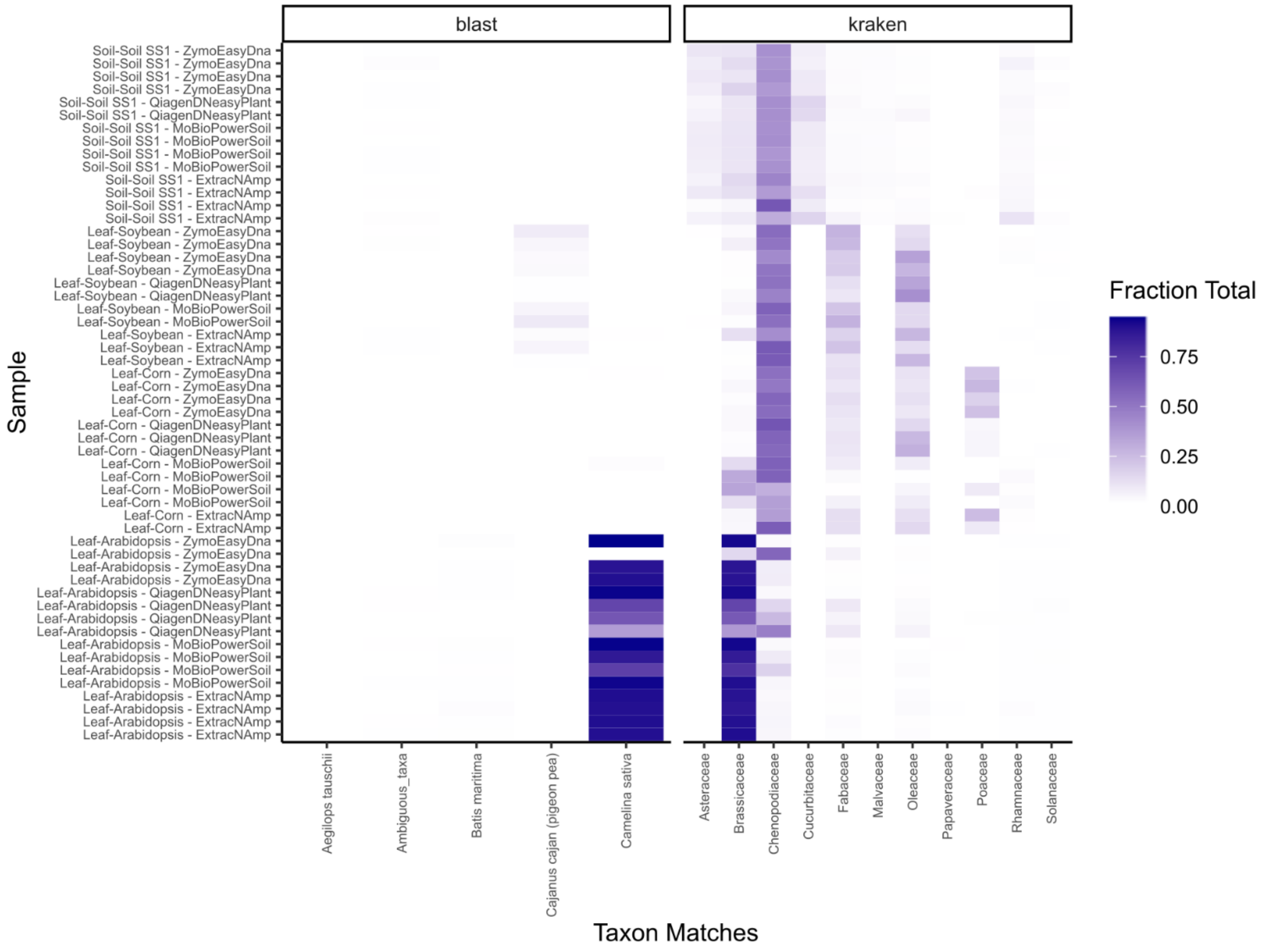


**Supplemental Figure S2 – Species check of plant extraction samples.** Due to the strong difference of maize leaf samples extracted by the PowerSoil method, chloroplast sequences in each plant extraction sample were checked for species identity by both BLAST ((Camacho et al. 2009); left) and Kraken2 ((Wood, Lu, and Langmead 2019); right). Low diversity of chloroplast sequences coupled with sequencing errors means that these programs often did not tag the correct species of origin. However, patterns of species identification are consistent within sample types. (For example, the Arabidopsis samples BLAST to *Camelina sativa* and kraken identifies them as part of the Brassicaceae.

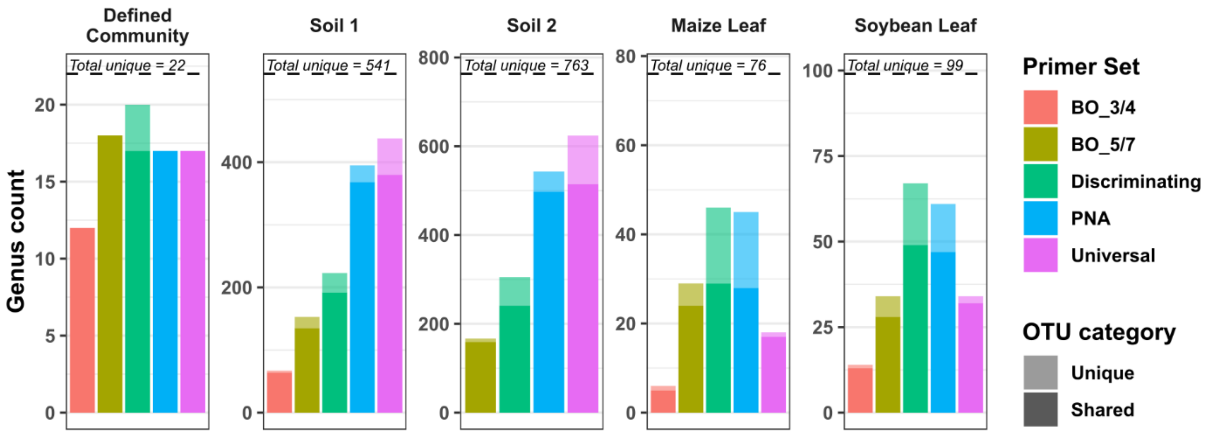


**Supplemental Figure S3 – Shared and unique genera among amplification methods.** The number of shared (dark) and unique (light) genera identified by each method is shown relative to the total number of unique genera (dotted line). A genus is counted as "shared" if at least two methods identified members of that genus.
